# Single-cell Pairwise Relationships Untangled by Composite Embedding model

**DOI:** 10.1101/2022.09.16.508327

**Authors:** Sishir Subedi, Yongjin P. Park

**Affiliations:** Bioinformatics Graduate Program, University of British Columbia, BC, Canada; BC Cancer Research, Part of Provincial Health Care Authority, BC, Canada; Department of Pathology and Laboratory Medicine, University of British Columbia, BC, Canada; Department of Statistics, University of British Columbia, BC, Canada

**Keywords:** Single-cell genomics, network, cellular communication, surface protein interaction, deep learning, variational autoencoder, probabilistic topic modelling

## Abstract

In multi-cellular organisms, cell identity and functions are primed and refined through interactions with other surrounding cells. Here, we propose a scalable machine learning method, termed SPRUCE, which is designed to systematically ascertain common cell-cell communication patterns embedded in single-cell RNA-seq data. We applied our approach to investigate tumour microenvironments consolidating multiple breast cancer data sets and found seven frequently-observed interaction signatures and underlying gene-gene interaction networks. Our results implicate that a part of tumour heterogeneity, especially within the same subtype, is better understood by differential interaction patterns rather than the static expression of known marker genes.

## Introduction

The advancement in single-cell RNA-sequencing (scRNA-seq) has emerged as a new frontier in genomics. Quantification of multimodal omics at a single-cell resolution has made it possible to gain insights into different aspects of cancer biology^1^. One of the fundamental questions in cancer research is how cancer cells interact with each other in a confined heterogeneous environment such as a tumour microenvironment (TME). Studies have shown that cell-cell communication (CCC) among cell populations in the TME is crucial in cancer growth and metastatic processes^2^. Understanding the intricacies of communication among tumour and their interacting partner cells could aid in identifying a potential therapeutic avenue in cancer.

A significant technical challenge that stands in understanding the dynamics of cell-cell interactions in TME is devising a systematic approach to isolate and capture interaction signals from each interacting cell pair. A conventional approach to studying CCC involves clustering features in low-dimensional space and inferring interactions between clusters of know cell types^3–5^. While these methods have uncovered numerous signalling mechanisms that govern cellular differentiation and pathogenesis, they assume each cluster, annotated using a limited number of marker genes, represents a cell type; hence all the cells within a cluster interact in the same manner. These methods do not account for intracluster cellular heterogeneity. Cells within a cell type may exist in multiple subtypes/states and manifest heterogeneous interaction patterns based on the type and state of the interacting partner cell, which is critical in understanding cancer progression^2,6^. Additionally, interaction among cells in different contexts, such as disease states, are studied separately, which loses context-specific variability information and are repetitive and computationally expensive.

Recent studies have addressed these challenges and developed methods to capture the diversity of cell interactions within the same cluster. Tensor-cell2cell^7^ uses tensor-based dimensionality reduction techniques to infer context-driven CCC pattern. scTensor^8^ also uses a tensor decomposition algorithm to infer many-to-many cell pair relationships as a hypergraph. These methods rely on a priori knowledge of cell type and aggregating cells to calculate communication scores based on the mean expression of ligand-receptor (LR) genes. SoptSC^9^ calculates signalling probability between two cells based on pathway-specific LR and target genes and addresses heterogeneity of cells within the same cluster. However, the method requires a user-defined comprehensive list of pathway genes and does not scale to cohort-level studies.

Here, we present a novel computational approach termed SPRUCE, Single-cell Pairwise Relationship Untangled by Composite Embedding, to analyze tens of millions of cell pairs in a scalable way. Adopting known ligand and receptor protein-protein interactions, we asked how and why cell pairs are localized in the proximity of latent topic space and highlighted common patterns repeatedly observed in tumour microenvironments data. SPRUCE is based on an embedded topic model (ETM), a generative deep learning method built on variational autoencoder architecture, and represents single-cell vector data in low-dimension topic space with an interpretable topic-specific gene expression dictionary matrix. It has been successfully implemented in natural language processing to extract meaningful topics representing large-scale documents^10^. A recent study, scETM, showed that ETM-based techniques efficiently capture essential biological signals from sparse and heterogeneous single-cell data^11^. The key contribution of our approach is the unbiased identification of interpretable cell subtype/state across multiple datasets by characterizing LR genes-driven patterns of cell-cell interactions. Existing graph-based single-cell analysis methods often define cell-cell interaction modules as densely-connected components in a graph (an adjacency matrix). Our SPRUCE model considers cell-cell interaction patterns as a stream of edges, or a giant incidence matrix (edge by vertex or other vertex property).

## Results

### Overview of SPRUCE model training in breast cancer study

We combined existing breast cancer data sets^12,13^ and cancer-specific immune cell data^14^ and constructed a comprehensive single-cell catalogue for an unbiased breast cancer study, yielding a data matrix consisting of 20,288 genes and 155,913 cells. We first mapped cells from multiple data sets onto a common latent topic space (K=50) and harmonized them based on variational autoencoder-based topic modelling, not posing additional assumptions, such as a selection bias imposed by top marker genes. Based on cosine similarity in the latent topic space between cells, we then constructed cell-cell interaction networks and performed stratified sampling so that topic-topic relationships are similarly represented in the subsequent steps in SPRUCE training. For each of these 25M+ cell pairs, we extracted gene expressions of the known 648 ligand and 672 receptor proteins^15^ and used them as feature vectors for SPRUCE model training.

### Multinomial probabilistic topic modelling identified 50 cell topics across 11 known cell types

We implemented a Bayesian deep learning approach to estimate embedded topic models across 155,913 cells with 50 latent dimensions (Fig. 1A). We found each cell topic corresponds to a group of an average of 3118 cells (with a standard deviation of ± 5,987) (Fig. 1B). Among 50 topics, 32 of them contained more than 100 cells. The highest number of cells (24% of the dataset) were assigned to topic 37, in which 96% of cells were previously identified immune cells (T and B cells). 98% of cancer cells from the dataset were assigned to 13 cell topics. The cancer cell proportion in 9 of 13 topics was greater than 95%. The latent cell topics with cell type annotated were visualized with UMAP, showing distinct clusters for each topic where the majority of cells belong to one of the major types of cells in the dataset (Fig. 1C). These clusters show a unique set of top genes associated with each topic identified using the optimized gene loading matrix from the model (Fig. 1D). We further confirmed concordance with the previous analysis conducted in original papers using cell type lineage canonical markers (Fig. 1F). The estimated cell topic proportions show that the resident cell types have similar topic proportions. However, cancer cells have a different mixture of topic proportions which shows that the model identified many distinct topics of cancer cells (Fig. 1E). We also tried a different number of cell topics from 10, 25, and 50 and decided to use the 50-topic model because major cell types, especially cancer cells, showed well-separated distinct clusters (Fig. S1A-B).

**Figure 1.**
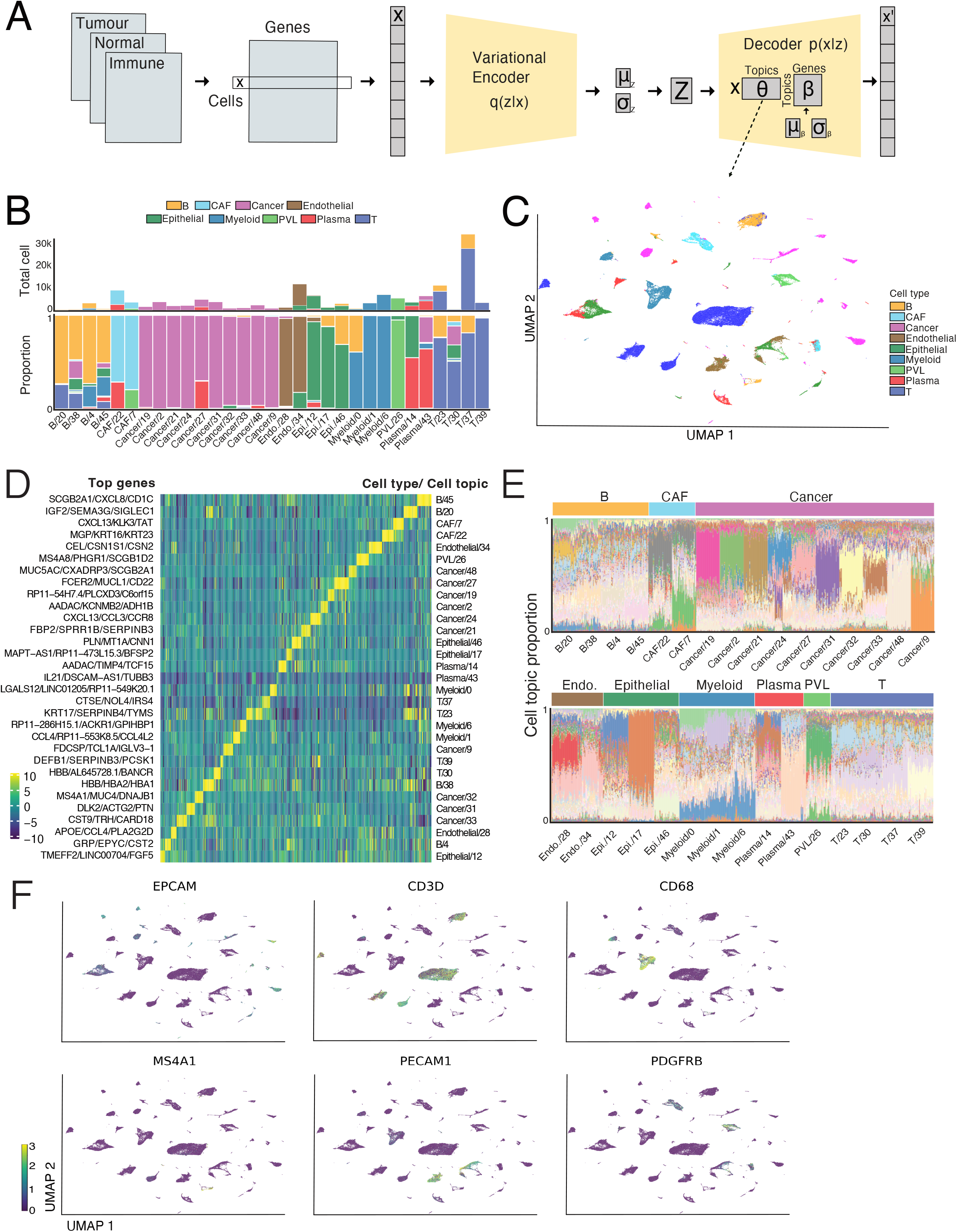
Probabilistic topic model identifies cell topics for resident cell types and cancer sub-types. (A) Given an integrated single-cell data from breast cancer tumour microenvironment, SPRUCE cell topic model learns the latent topic representation of the cells and generates biologically interpretable topic embeddings separately over the genes. (B) Distribution of cell types highlighting the total number of cells in each cell topic. Relative proportion of cell types in each cell topic. Cell topic is assigned to each cell based on the highest score. (C) UMAP visualization of 50 cell topics representing an integrated dataset consisting of 155,913 cells. Each dot is a single cell, and colors represent the corresponding cell type clustered according to k-means(n=50) algorithm on cell topic proportions and annotated using the majority voting rule with annotation from previous studies and SingleR. (D) Heatmap of top 25 gene loadings associated with each cell topic with the total number of cells > 100. Cell topics (y axis) and genes (x axis) are ordered according to hierarchical clustering (optimal leaf ordering). The cell type and topic are shown on the right y-axis, while the top 3 genes for each cell topic are depicted on the left y-axis. (E) Relative proportion of cell topics from a sample(n=50 cells per topic) of cells assigned to cell type - cell topic pairs. (F) Log normalized (total count to 10,000 reads per cell) expression of markers genes for cell types-*EPCAM* for epithelial, *CD3D* for T-cells, *CD68* for myeloid cells, *MS4A1* for B-cells, *PECAM1* for endothelial cells, and *PDGFRB* for CAF/PVL cells.

### Seven robust TME-specific interaction signatures were found in 25 million cell-cell pairs

We constructed LR gene expression data from 155,913 cells to construct a set of 24,790,167 cell pairs and estimated embedded interaction topic models with 25 latent dimensions (Fig. 2A-B). The interaction topic model recapitulates two types of cell-cell communication mechanisms. First, the model captures interactions between differentially expressed ligand and receptor genes in different cell types. Second, the model considers ligand and receptor genes correlated within each cell type but may not be differentially expressed. Among 25 interaction topics representing 25 million cell pairs, seven topics (2, 4, 7, 10, 18, 22, and 24) represented 55% of the total cell pair interactions, with each topic containing 3%+ cell pairs (Fig. 2C). The other 18 interaction topics, each with 2% of the total cell pair interactions embedded baseline interaction signal. The most represented interaction topic was topic 22, consisting of 12% of the total cell pair interactions.

**Figure 2.**
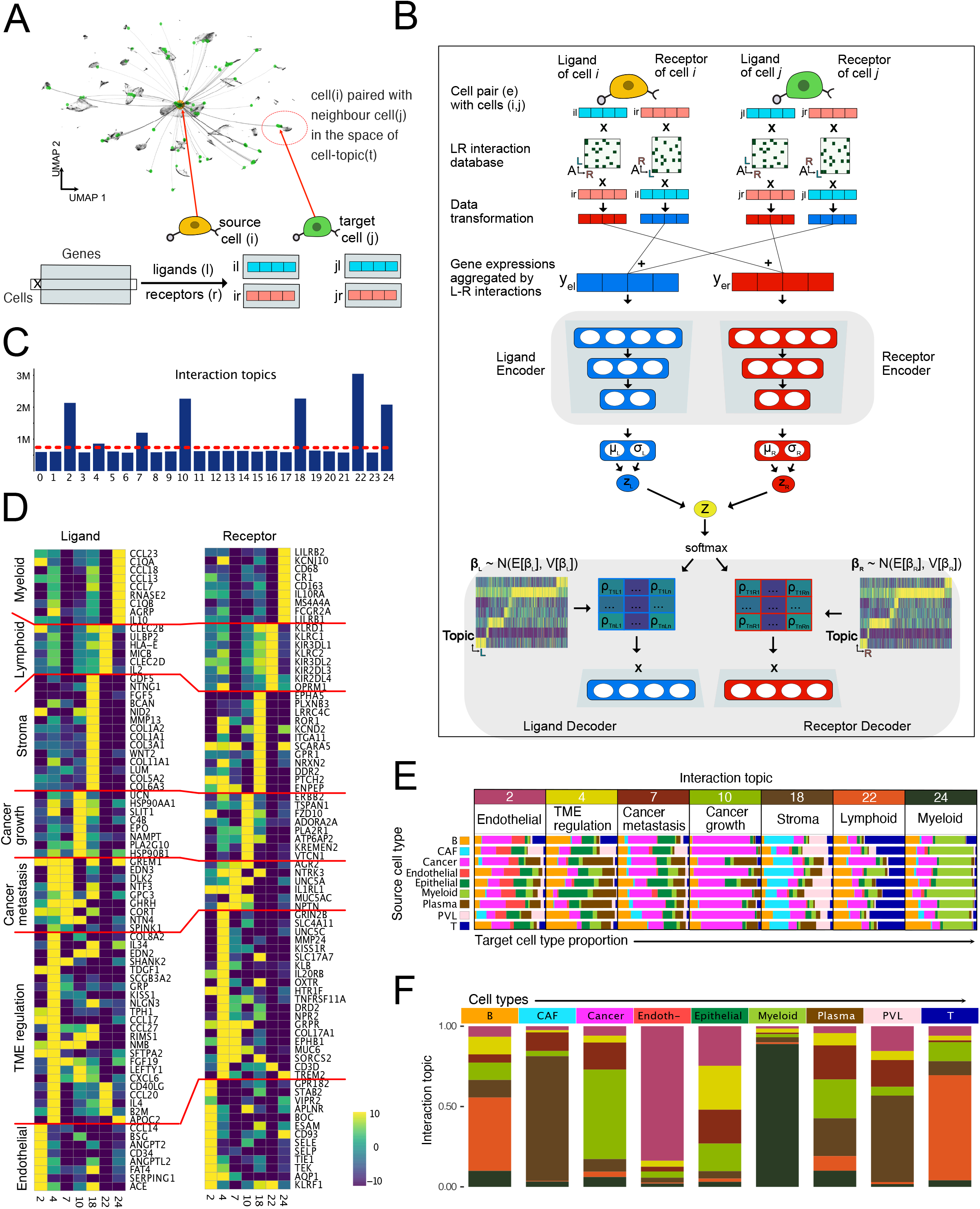
SPRUCE model overview and the common interaction patterns identified by the model. (A) UMAP-based representation of cell pairing method where source cell is paired with four different target cells from each cell topic. (B) Given a cell pair LR gene expression data as input, SPRUCE model transforms and aggregates interaction-driven LR data space and feeds into the neural network. The encoder learns the latent topic representation of the interaction between cells using the mixture of experts approach from ligand and receptor encoders. The decoder generates biologically interpretable topic embeddings separately over the ligand and receptor genes. (C) The distribution of ∼25 million cell pair interactions over 25 interaction topics shows seven major interaction patterns. (D) Heatmap of top 10 gene loadings associated with seven interaction topics. Interaction topics (x axis) and LR genes (y axis) are ordered according to hierarchical clustering (optimal leaf ordering). (E) Relative proportion of target cell type in all the source cell types associated with the major interaction patterns. Each row is source cell type (y axis) and target cell type proportions (x-axis). (F) Enrichment of interaction patterns among cell types.

The model estimated the LR gene loadings in each interaction topic that described the relative contribution of each gene. These loadings can be ranked to identify biologically interpretable topic-specific top genes in each interaction topic (Fig. 2D). Topics 22 and 24 captured immune-related interactions. Topic 22 was labelled as a lymphoid topic because top receptor genes included the known subunit of T-Cell Receptor Complex *CD3D* and killer cell lectin-like receptors *KLRC1, KLRC2*, and *KLRD1*. The top ligands in this topic are *HLA-E, CLEC2B*, and *CLEC2DC*, which are essential known modulators in cytotoxic T cells^16^. Similarly, topic 24 was labelled as a myeloid topic as top genes in this topic showed enrichment of LR genes expressed by myeloid progenitors, for example - receptors such as *CD68, TREM2*, and *CR1*, and ligands such as *CCL23, CCL18, CCL13*, and *C1QA*^17^.

Topics 10 and 7 represented many oncogenes mutated in cancer. For instance, Topic 10 was cancer-growth associated, and genes that play a role in cancer cell survival and growth are enriched in this topic. The top receptors in this topic are growth factor receptors such as *ERBB2*, cell proliferation and growth signalling receptor *FZD10*, and immune inhibiting signalling receptor *ADORA2A*^18,19^. Similarly, Topic 7 was a cancer-metastasis topic and genes such as *NTRK3*, known to increase the metastatic potential of cancer cells, *GRPR*, which promotes EMT, and *UNC5A*, a known regulator of cancer plasticity, are enriched in this topic^20–22^.

Moreover, Topic 18 was stroma-specific and represented genes that play an integral role in regulating the extracellular matrix (ECM) of the tumour immune microenvironment. These genes are highly expressed in cancer-associated fibroblast (CAF) and perivascular-like (PVL) cells. The top ligands in these topics are *COL1A1, COL1A2, COL3A1*, and *MMP13*, and the top receptors are *ITGA11* and *SCARA5*^23,24^. Similarly, Topic 2 is enriched with genes highly expressed in endothelial cells, thus labeled an endothelial topic. Here, ligand proteins highly expressed in endothelial cells, such as *CD34, ANGPT2*, and *NID2* and receptors such as *APLNR* and *ESAM* are enriched^25^. Likewise, Topic 4 enriches genes involved in TME regulation processes and is thought to facilitate cancer progression and growth. The top genes in this topic include *KISS1R*/*KISS1*, which play a complex role in both restricting and promoting cancer cell survival, *IL20RB*, which promotes immunosuppressive microenvironment, and *MMP24*, which negatively regulates the aggressiveness of cancer cells^26^.

The top LR genes in the major interaction topics show enrichment of different cell type-specific functional interactions. To confirm that each interaction topic captured cell type-specific CCC, we took a closer look at the distribution of cell types of target cells in each interaction topic for all the cells in the dataset. We found that the functional role of enriched top LR genes in each interaction topic matched with the dominant cell type of target cells in that topic (Fig. 2E). For example, in the cancer-growth associated, on average, 68% of target cells for all cell types were cancer cells. Similarly, 49% of target cells in the stroma topic were CAF/PVL cells, and myeloid and T cells comprised 49% and 38% of target cells in the myeloid and lymphoid topics, respectively. For the endothelial topic, the dominant target cell type stemmed from endothelial cells, comprising 18%. In contrast, for the TME-regulation topic, both epithelial and plasms cells were dominant cells consisting of 20% and 22%, respectively.

In addition, the cell type-specific enrichment of interaction topics was further corroborated by the distribution of interaction topics in each cell type. Cancer cells, along with epithelial, plasma, and B cells, showed heterogeneous interaction patterns compared to myeloid, T, endothelial, CAF, and PVL cell types (Fig. 2F). For cancer cells, the majority of interactions belonged to the cancer-growth and cancer-metastasis topics, where many of the top genes were oncogenes. Here, 55% of the total interactions were incident with a cancer-growth process, and 17% belonged to the cancer-metastasis topic, while the other five remaining topics consisted of 3-8% of interactions. Among the non-malignant cell types, the dominant interaction topic for myeloid cells was the myeloid interaction topic, and for T-cells, it was the lymphoid interaction topic. Here, 88% of myeloid cell and 65% of T cell interactions were found to be in respective interaction topics. Similarly, 83% of cell interactions with endothelial cells belonged to the endothelial topic, and 77% and 53% of interactions with CAF and PVL cells, respectively, were the stroma topic. In contrast, plasma, B, and epithelial cells showed a higher mixture of non-immune associated interaction topics.

### Interaction topics provide a way to understand breast cancer heterogeneity

Next, we investigated the heterogeneity of breast cancer cells based on the unbiased transcriptomic signature captured by the cell topic model while relating the cell topics to the interaction topics characterized by the SPRUCE analysis. The interaction patterns of 25,835 cancer cells manifested all the patterns of interactions (Fig. 3B). The cell topic model identified 13 cell topics for cancer cells that show a distinct pattern of interactions with their target cells (Fig. 3C). As expected, the cancer growth topic was the most dominant (>57%) among 7 of 13 cell topics. For instance, 75%, 70%, and 68% of interactions for cancer cells in cell topics 24, 48, and 2 were driven by cancer-growth-related interactions. However, two cell topics were found more frequently interacting with non-cancer cells. The 66% of interactions involving (cell) Topic 9 were of stroma edges, and 60% of edges emanating from Topic 34 were assigned to endothelial interactions. Such a striking interaction heterogeneity was rarely observed in other cell types, such as T-cells, myeloid, endothelial, CAF, and PVL cell types, as they mostly interact within the same cell types. B-cells and epithelial cells, however, showed substantial variability of interaction patterns across cell topics.

**Figure 3.**
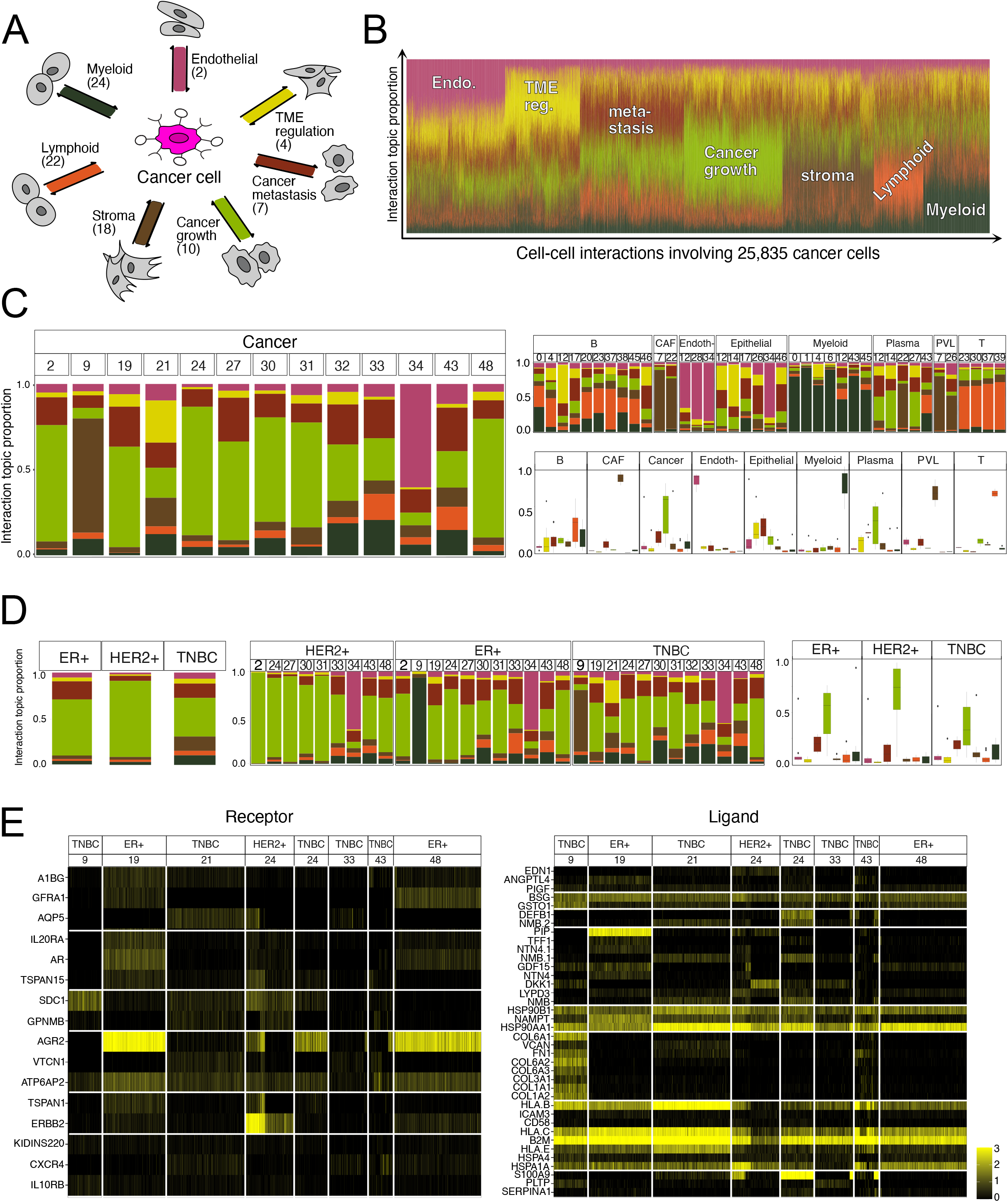
Heterogeneity of cancer cells revealed by the cell-topic specific interaction patterns. (A) Representation of interaction topics of cancer cells with surrounding cell types in the TME. (B) Structure plot showing the variability of interaction topic proportion estimates among 25,835 cancer cells in the dataset. Proportion of seven interaction topics (y axis) and cancer cell pair (x-axis) with one representative target cell among 159 source-target cell pairs are clustered using k-means(n=7) clustering algorithm using interaction topic proportions. (C) The proportion of interaction topics of cancer and non-malignant cells associated with cell topics. Boxplots show the distribution of interaction topic proportions for each interaction topic across all cell topics. (D) The proportion of interaction topics of cancer subtypes associated with cell topics. Boxplots depict the estimated interaction topic proportions across all cell topics. (E) Log normalized (total count to 10,000 reads per cell) expression of LR genes from all the cells associated with respective cancer subtype and cell topic. The genes (y axis) are top 25 LR genes from seven interaction topics with expression values > 0.05.

Breast cancer cells are classified into subtypes based on the genomics and pathology of the disease, and different subtypes often result in markedly different clinical outcomes^27^. All three different subtypes of cancer cells indeed exhibit diverse interaction patterns where TNBC (triple-negative breast cancer) cells were more heterogeneous compared to HER2+ and ER+ subtypes (Fig. 3D). Here, more than 85% of interactions of the HER2+ subtype consisted of cancer-related topics, such as 82% for the cancer-growth and 5% for cancer metastasis topics. Similarly, for the ER+ subtype, more than 80% of interactions were cancer-related: 61% for the cancer-growth and 20% for the cancer-metastasis topics. In contrast, TNBC subtype cells were more diverse in interactions, with 42% for cancer growth, 15% for the cancer-metastasis, 15% for stroma, and 10% for myeloid topics.

Additionally, our approach identified a specific group of cells (cell topics) within these cancer subtypes that show topic-specific interaction patterns. For instance, TNBC cell topics show higher heterogeneity in interaction patterns at the cell topic level compared to the ER+ and HER2+ subtypes. The distribution of interaction patterns among cell topics is correlated with the expression pattern of LR genes enriched in each interaction topic. For example, TNBC cancer cells in cell topic 9 show higher expression of LR genes enriched in the myeloid interaction topic. In contrast, the cancer-growth LR genes are dominant among TNBC cancer cells in cell topic 24 (Fig. 3E).

### Interaction topics uncover underlying cancer-subtype-specific gene-gene interaction networks

Our approach also reveals topic-specific interaction patterns in the model parameter matrix (Fig. 4), with which gene-gene correlation networks can be estimated (ligand vs. receptor). For instance, gene networks in the cancer metastasis topic (Fig. 4A, burgundy strips; Fig. 4B, the first panel) consisted of a group of structural genes *CLDN4, LSR*, and *DSG2* involved in cell transformation and migration and signalling pathways *GPR37, CD151*, and *CD63* that are active in proliferation and migration, including epithelial-mesenchymal transition^28,29^. In the cancer growth topic (Fig 4A, green strips; Fig. 4B, the second panel), the resulting gene network primarily consisted of known oncogenes, such as *PTPRF, FGFR1, ERBB2*, and *TNFRSF1A*^30^ and developmental genes, such as *LAMP1, ITGB1, RPSA, CANX, ATP6AP2*, and *MCFD2*^31^. It is worth noting that the interaction networks adjacent to cancer cells generally include known cancer-related genes and other genes involved in cellular developmental process. Our analysis put them together in the same network modules, implicating a potential role of these interactions for cancer cells to hijack a normal cellular process. Similarly, immune-modulatory receptors primarily expressed in myeloid lineage cells (Fig. 4A, dark green; Fig 4B, the third panel) *TREM2, CSF1R, CSF2R, LILR*, and *IL3R* and signalling pathways *LTBR* and *TYROBP* required for the activation of myeloid cells are associated with myeloid-associated interaction topic. This topic captures the interactions between cancer cells and myeloid cell progenitors such as tumour-associated macrophages (TAM) in the tumour microenvironment, suppressing T cells and facilitating tumour growth^32^.

**Figure 4.**
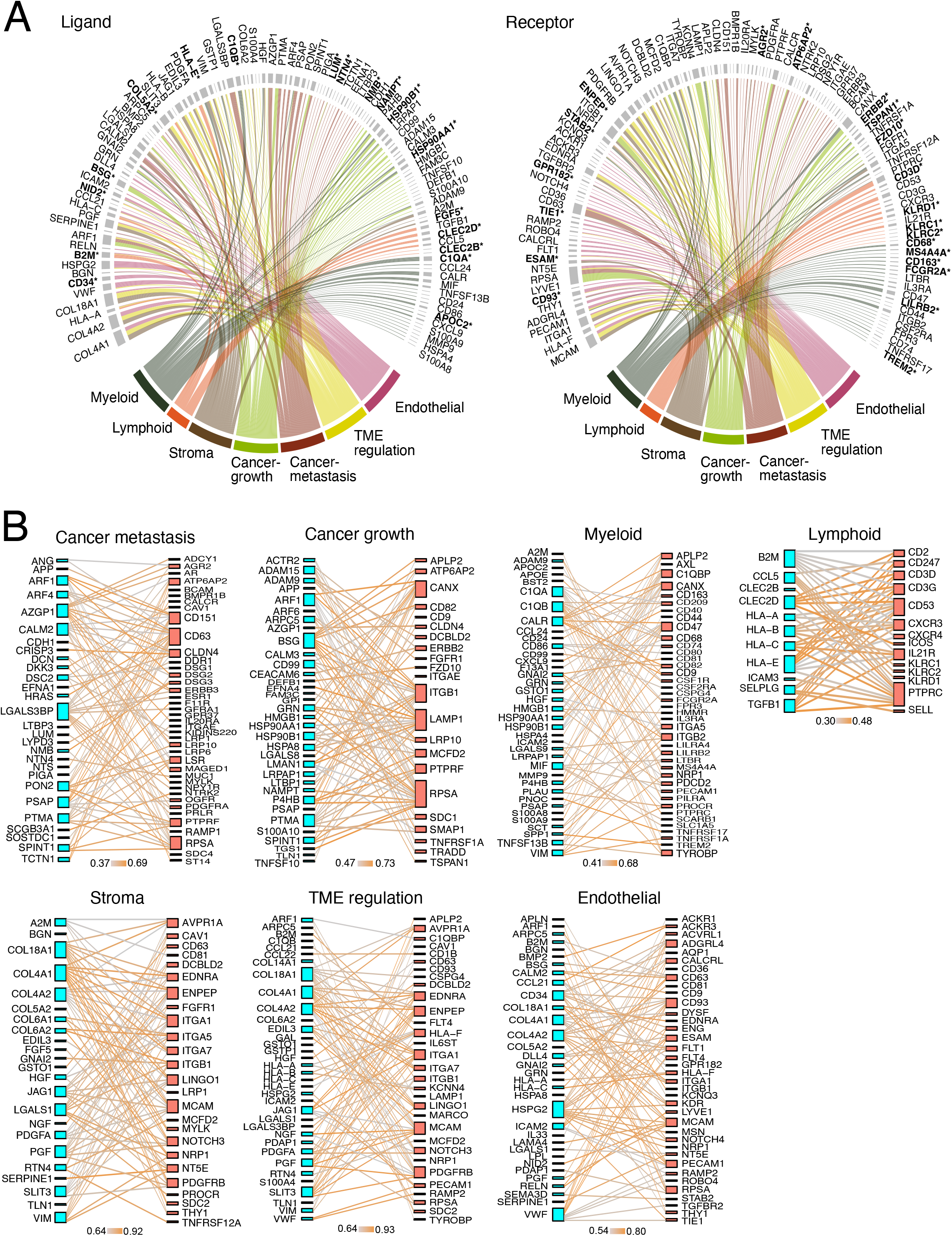
Gene network derived from the interaction topics. (A) Circular chord diagram of the interaction topic-gene network representing significantly (Z-score > 4.0) enriched LR gene loadings in each interaction topic. The edge between LR genes and interaction topics shows the occurrence of a gene in LR pairing and if a gene is unique to or common among different interaction topics. The genes with (*) symbol are among the top 10 LR genes based on loadings estimated by the interaction topic model. (B) Ligand-receptor bipartite network for each interaction topics depicting significantly (Z-score > 4.0) enriched LR genes. The edge in the graphs represents the magnitude of Pearson correlation coefficients. The top 100 LR edges in each interaction topic with a correlation coefficient > 3.0 are selected.

In the lymphoid topic (Fig. 4B, the fourth panel), T cell receptor (TCR) complex (*CD3D, CD3G, CD2*, and *CD247*) and TCR signalling pathway genes (*PTPRC, CD45*, and *CD53*) are highly expressed both in cancer and surrounding immune cells^33^. Other chemokine receptor genes, such as *CXCR3* and *CXCR4*, were also found highly co-activated in this interaction topic, corroborating the pivotal role of crosstalk between T cells and cancer cells in promoting cancer growth, immune evasion, and metastasis^34^. Interestingly, other killer-cell lectin-like receptors, which also co-occurred in this module, namely *KLRC1, KLRD1*, and *KLRF1*, are known to restrict T-cell’s antitumour immunity^35^. The stroma topic (Fig. 4A, brown strips; Fig. 4B, the first panel of the second row) represented gene networks that capture the interaction of cancer cells with surrounding cells that promote its vascularization. It consisted of *NOTCH3, AVPR1A, MYLK* and integrin-mediated *ITGA1, ITGA5, ITGA7*, and *ITGB1* signalling pathways that play vital roles in tumour cell adhesion and progression^36^. The genes *MCAM, ENPEP, EDNRA*, and *DCBLD2* that promote blood vessel formation and enhance tumorigenesis are enriched in this topic^37^. Similarly, genes enriched in the endothelial topic recapitulate interactions of cancer cells in developing tumour vascular networks, especially in conjunction with endothelial cells. Previously known that *PECAM1, CALCR, ADGRL4*, and *CD93* genes are predominantly expressed in endothelial cells and regulate angiogenesis in tumour cells^38^. Additionally, TME-regulation associated topic primarily consisted of genes mixture of endothelial-associated and stroma-associated topics with enrichment of distinct genes known to control tumour growth and promote stemness of cancer cells in microenvironment such as *KCNN4, IL6ST*, and *CD1B*^39^. Further, we observed a significant overlap between the gene-gene interactions derived from the interaction topic model and the gene network in the STRING database^40^. Here, 51%, 40%, 18%, 25%, 13%, 12%, and 13% of ligand-receptor pairs (Z-score >4.0 and Pearson correlation coefficient > 3.0) matched with gene pairs in the STRING database (combined score > 0.3) from lymphoid, myeloid, stroma, endothelial, TME regulation, cancer metastasis, and cancer growth topics, respectively (Fig. S3).

## Discussion

No single cell can exist alone in human tissues. We propose a novel machine learning framework that systematically dissects tens of millions of cell-cell pairs and uncovers common patterns of how cells talk to each other concerning cell surface ligand-receptor protein interactions. In particular, our analysis focused on finding commonly used communication channels in breast cancer progression and metastasis by reanalyzing state-of-the-art single-cell genomics data sets. Our approach, built on probabilistic topic modelling and variational autoencoder model, specifically demonstrated that a part of cancer heterogeneity could be understood in diverse and context-specific interaction partners of cancer cells. We found that many ligand-receptor interactions can occur in a subtype-specific manner, although cells are largely clustered as cancer cells in conventional single-cell analysis. Along the line, our results suggest that cell types and states are better understood, and the definitions of cell types can be refined while considering cell-cell communication patterns.

Our proposed approach generalizes existing bioinformatics methods and does not rely on prescribed cell-type annotations/clustering results, which may introduce unwanted biases in downstream analysis. Moreover, if cells within a cluster are not homogeneous as anticipated, a clustering-based cell-cell commutation method can easily result in confounded correlation statistics, clearly violating necessary assumptions, such as independent and identification distributed expression values. Here, we differently formulate cell-cell communications analysis from an edge’s perspective, whereby a single edge (interaction) is a data point, and the feature vector can be engineered by exploiting the information of both endpoints of the edge (ligand and rector expression values). Such a novel formulation is better suited for the analysis of large-scale single-cell data and also easily extends to principled data integration strategies. For instance, if cell-cell interaction pairs were already constructed by spatial transcriptomics data, we can easily construct feature vectors by combining two gene expression vectors (one from the source and the other from target cells). For the multiomics data integration tasks, we can concatenate multiple data modalities to investigate the co-occurrence of multimodal expressions, such as DNA accessibility, histone modifications, and metabolomics.

## Limitation of the Study

We acknowledge that our SPRUCE approach relies on several specialized modelling assumptions. One of which is that we assume that known ligand-receptor protein-protein interaction networks serve as a superset/backbone of topic-specific interaction networks. Considering that most protein-protein interactions were experimentally discovered *in vitro* by error-prone high-throughput methods, there is room for improvements in terms of the precision and specificity of the interaction analysis method. Here, we only focused on immediate surface protein interactions. However, establishing causal effects on the downstream genes emanating from surface protein singling pathways can further enrich our understanding of disease etiology.

## Data and Code Availability

## Acknowledgments

This work was supported by the BC Cancer Foundation and NSERC Discovery Grant (YPP). We acknowledge financial support from UBC Four Year Fellowship (SS). Much of the computation was supported by Cascadia Data Alliance Award (title: Eavesdropping Communications between Cancer and Immune Cells).

## Author contributions

Conceptualization, Y.P.P.; Methodology, Y.P.P. and S.S.; Investigation, S.S.; Writing – Original Draft, S.S. and Y.P.P.; Writing – Review & Editing, S.S. and Y.P.P.; Funding Acquisition, Y.P.P.; Resources, Y.P.P; Supervision, Y.P.P.

## STAR Methods

### RESOURCE AVAILABILITY

#### Material availability

We made our source code and data sets available in the public repository:

#### Lead contact

Should the readers have trouble accessing data and source code, the lead and corresponding authors can be reached to facilitate data and knowledge transfer.

### EXPERIMENTAL MODEL AND SUBJECT DETAILS

We constructed a dataset to represent an immune-enriched breast cancer microenvironment by combining cancer cells with immune cells and healthy cells from three recent breast cancer-related studies. The breast cancer dataset consists of 100k cells from a single-cell atlas of human breast cancers^12^. The source of the normal dataset consisting 48k cells is a single-cell atlas of the healthy breast tissues^13^. The third dataset composed of immune cells is 6k subset of breast cancer CD4 and CD8 T-cells from a pan-cancer atlas of tumour-infiltrating T cells profiled across 21 cancer types and 316 donors^14^. The total number of cells in the combined dataset is 155,913. We filtered out genes detected in less than three cells along with mitochondrial and spike genes, leading to 20,265 genes in the final dataset.

### METHOD DETAILS

#### Topic modelling

SPRUCE takes a topic modelling approach to identify cell subtypes/states and define their interaction pattern based on cell-cell communication in the tumour microenvironment (TME). The model consists of two types of autoencoder-based topic models, each with a pair of encoder-decoder networks. The first model takes gene expression single-cell data as input and models cell topic and topic-specific gene loadings. Next, we assigned a cell topic to each cell based on the highest topic proportion from the cell topic model. The topic assignment was used to construct a set of target cells for each cell such that one cell is paired with the five nearest target cells from each topic. The LR gene expression data from each cell pair were transformed into each other’s space by a binary cell-interaction database– `CellTalkDB`. The transformed ligand and receptor data were treated as two independent modules with separate encoder and decoder modules in the model. The latent variables with encoded information from two modules were combined by taking their average to obtain a final interaction topic variables as a mixture of experts^41^ from the ligand and receptor latent space. The second ETM, the interaction topic model, uses transformed LR data from source-target cell pairs as input and models interaction topic for each cell pair with topic-specific LR gene loadings.

#### Notations

The following notation will be used to describe the data and model. Notations for gene expression and interaction data:

- *i* ∈ [*N*]: an integer index for a cell *i* of total *N* cells
- *g* ∈ [*G*]: an integer index for a gene *g* of total *G* genes
- *X*_*ig*_: gene expression (non-negative) count data measured on a gene *g* in a cell *i*

Notations for the cell topic model:

- *k, t* ∈ [*K*]: an index for a topic *t* of total *K* topics
- *θ*_*it*_ : cell topic proportion of *i*th cell for *t*th topic (*θ* _*it*_ > 0 and 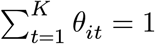 for all *i*)
- *β*_*tg*_(cell topic model): gene proportion of *t*th cell topic for *g*th gene

Notations for the interaction topic model:

- *e*: an index for a cell pair, e.g., *e* = (*i, j*) for a cell *i* and *j*.
- *X*_*li*_: expression count of a ligand protein *l* in a cell *i*
- *X*_*ri*_: expression count of a receptor protein *r* in a cell *i*
- *Y*_*el*_: transformed and aggregated count data of a ligand protein *l* in a cell pair *e*
- *Y*_*er*_: transformed and aggregated count data of a receptor protein *r* in a cell pair *e*
- *θ*_*et*_: interaction topic proportion of *e*th cell pair for *t*th topic
- *β*_*tg*_ (interaction topic model): gene proportion of *t*th interaction topic for a gene *g*, which can be either a ligand protein/gene *l* or a receptor protein/gene *r*.

#### Cell-level probabilistic topic modelling

Firstly, we designed a topic model for cell type annotations, treating cells as documents and genes as vocabulary, built on the Embedded Topic Model framework^10^. ETM generally outperforms traditional topic modelling approaches, such as Latent Dirichlet Allocation^42^, relying on tailored variational inference^42^ and collapsed Gibbs sampling inference^43^, and fits naturally in a variation autoencoder (VAE) framework^44^ while providing a scalable GPU-based inference algorithm.

Letting **x**_*i*_ = (*X*_*i*1_, …, *X*_*iG*_) be a vector of gene expression counts on *G* genes for each cell *i*, a topic modelling assumes that **x**_*i*_ were generated by multinomial distribution parameterized by a normalized gene expression frequency vector, namely *ρ*_*i*_ = (*ρ*_*i*1_, …, *ρ*_*iG*_), achieving a scale-invariant property across different cells, batches, and data sets:

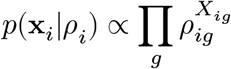

In the original ETM formulation^10^, *ρ* is directly modelled by transforming each cell’s topic proportion *θ*_*it*_ in a topic space to a gene space as a linear combination of topic-specific probabilities, *ρ*_*ig*_ = ∑_*t*_ *θ*_*it*_*β*_*tg*_, where *β*_*tg*_ captures a topic *t* specific frequency of a gene *g*. The latent cell topic proportion *θ*_*it*_ is drawn from Logistic Normal distribution with an auxiliary Gaussian vector **z**_*i*_ = (*Z*_*i*1_, …, *Z*_*iK*_):

Assuming **z**_*i*_ ∼ 𝒩(0, *I*) *a priori*, the encoder network will first generate

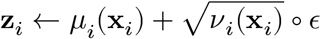

where the mean *μ* and variance *v* functions were modelled by deep neural networks, taking expression data **x**_*i*_, and the stochastic vector were simply generated by 𝒩(0, 1) independently. We can then project the Gaussian latent states into the desired topic space:

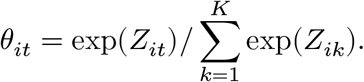

Since a Gaussian random variable can be generated by a reparameterization trick, which then separates model parameters from stochastic variables, the latent variables *θ* and model parameters *β* are seamlessly integrated in a neural network model; they can be optimized by back-propagation algorithm^45^ implemented in PyTorch library (https://pytorch.org/).

#### Bayesian autoencoder model for topic modelling

Instead of directly modelling *ρ*, we introduce the Dirichlet prior on the gene frequency and parameterize the Dirichlet as a generalized linear model(GLM) with linear combinations of topic-specific probabilities:

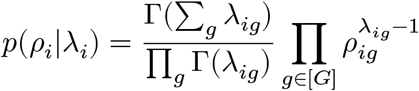

where Γ(·) is the Euler’s gamma function,

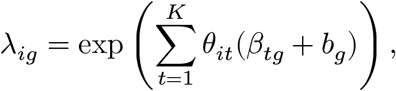

and

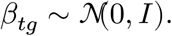

Exploiting the conjugacy between the multinomial and Dirichlet distributions, we can integrate out the unknown parameters *ρ*. Then, the marginal likelihood of a single cell data **x**_*i*_ then becomes:

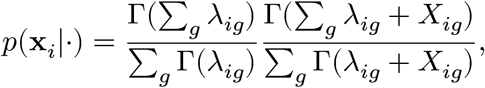

where Γ(·) is the Euler’s gamma function.

#### Variational inference algorithm

We resolved the posterior distribution of latent variables and model parameters, *p*({*θ*_*i*_}, {*β*_*gt*_}|{**x**_*i*_}), by finding variational/approximating distributions *q*(*θ*_*i*_|*μ*(**x**_*i*_), *v*(**x**_*i*_)) and *q*(*β*_*tg*_). We defined *q*(*θ*|·) as before using deep neural networks for Logistic Normal distributions. For the topic-specific gene matrix, *β*_*tg*_, we used mean-field approximation: 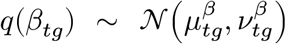. To minimize the Kullback-Leibler (KL) divergence between the true posterior and the approximate posterior, we maximize the evidence lower bound (ELBO) of the log-likelihood ℒ:

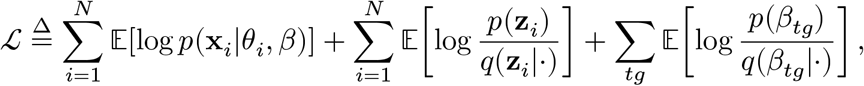

where the expectations were taken with respect to the variational distribution. The expectation operators can be well-approximated by summing over different **z**_*i*_ and *β* values sampled by the reparameterization tricks^44^.

The encoder for the cell topic model consisted of a 4-layered neural network with two hidden layers of size 200 and two with sizes 100 and 50. We used an Adam optimizer^46^ with a learning rate 0.01 and optimized the model for convergence for 1000 epochs with a minibatch size of 128 (Fig. S1D). The cell topics are almost ubiquitously present across cells from different data sets (Fig. S1C-E).

#### Cell type labelling by propagating within topic clusters

We generated a reference cell-type label for each cell in the dataset by combining the annotations from previous studies and cell type predictions from single-cell annotation tools. The cell annotations for breast cancer cells and immune cells from pan-cancer dataset were obtained from the previous studies^12,14^. For the annotation of normal breast cells, we used the reference-based cell type identification method `SingleR` with a tumour microenvironment reference dataset from CHETAH^47,48^. Next, we applied our proposed cell-topic model to an integrated dataset consisting of 155,913 cells and 20,265 genes. After unsupervised training of the model, all cells were mapped in the reduced latent cell topic space. We performed clustering on the reduced latent dimension using *k*-means algorithm (with *k* matched to the number of topics in cell topic model). Then each cluster was mapped to cell type using the majority rule on the reference cell type labels of cells assigned to the respective cluster, followed by relabeling of cells if needed for the downstream analyses.

#### Cell-cell interaction topic modelling

##### Construction of feature vectors for cell-cell interaction analysis

We assigned a cell topic to each cell in the dataset based on the highest proportion value from a vector of topic proportions inferred by the trained model i.e. topic assignment *t*_*i*_ for cell *i* is arg max(*θ*_*i*1_, *θ*_*i*2_, …, *θ*_*iK*_) where *θ*_*it*_ is cell topic proportion of *i*th cell for *t*th topic. Next, a set of target cells was constructed for each cell using the topic assignment from the cell topic model. For each cell, the five closest target cells from each topic were calculated using python package^49^ with angular distance on the cell topic space. An annoy model was created for each topic with a total cell count greater than 100 (32 out of 50 cell topics). We generated 159 target cell pairs for each cell - 32 topics and 5 target cells from each topic, excluding a self-target cell. Finally, a cell pair data matrix was constructed with 155,913 source cells times 159 target cells, consisting of 24,790,167 unique cell pairs.

The raw LR gene expression data from each cell pair is transformed into each other’s space using a binary ligand-receptor interaction matrix generated from a publicly available database–`CellTalkDB`^15^. Here, let *A* be a *l* x *r* binary matrix with its entries as *A*_*lr*_ = 1 if and only if a ligand *l* binds with a receptor protein *r* in the cell interaction database; otherwise, *A*_*lr*_ = 0. For each cell pair *e* ≡ (*i, j*), we aggregated expression counts for ligand and receptor proteins included in the *A* matrix by the reciprocal cell-cell interactions between *i* and *j*. For the activity for a receptor *r* ∈ [*R*], we combined expression values emanating from relevant ligand proteins/genes:

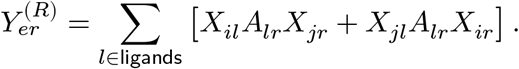

Similarly, for a ligand *l* ∈ *L*, we aggregated the values on the receptor side:

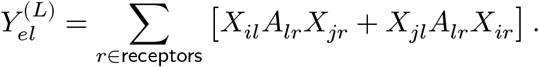

##### Interaction topic model

The interaction topic model uses transformed ligand and receptor (LR) gene expression data from source-target cell pairs to model the interaction topic for each cell pair and identify LR genes enriched in each interaction topic. The model follows the same architecture of the cell topic model, where each cell pair is considered a document and LR genes in the cell pair as words. We define the joint likelihood of the interaction topic model as product of two likelihood functions, each derived from the ligand and receptor topic models:

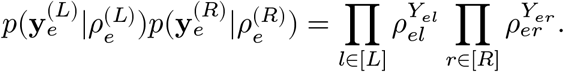

The decoder (data-generating) model for each side was defined as the same Multinomial-Dirichlet hierarchical model as the cell topic model. In order to map the ligand and receptor activities to a shared topic space, we formed a mixture of experts^41^ by equally mixing the outputs of two encoder networks, henceforth generating two sides of data with the same topic proportions. For each cell pair *e* ∈ [*m*], we generate the ligand and receptor activities and optimize the model parameters by the back-propagation in the following steps:

- Combine 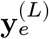 using **x**_*i*_ and **x**_*j*_ for ligand genes and 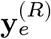 for receptor genes
- Generate latent state parameters, the mean 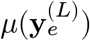 and variance 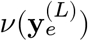 based on the ligand feature activities and the mean 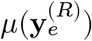 and variance 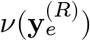 based on the receptor features
- Sample 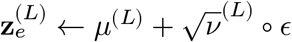 for the ligand and 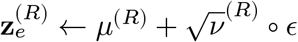 receptor networks
- Transform them into a common topic proportion vector 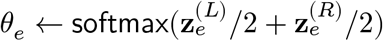
- Sample 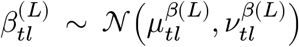 for ligand topics and 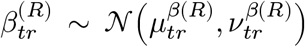 for receptor topics
- Compute the following composite ELBO objective ℒ^*LR*^, take stochastic gradients, and optimize by the Adam optimizer.

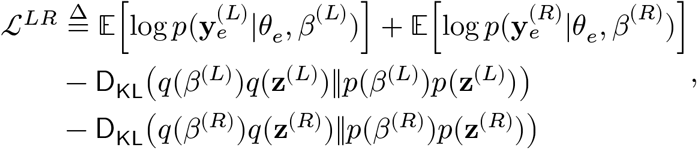

where we denote Kullback-Leibler divergence between *q* and *p*, i.e., 𝔼_*q*_[log *q*/*p*], by `D`_KL_(*q*‖*p*).

The encoder for the cell topic model consisted of a 3-layered neural network with hidden layers of sizes 200, 100, and 25. We used an Adam optimizer with a learning rate of 0.01 and optimized the model for convergence for 500 epochs with a minibatch size of 5088 cell pairs (32 individual cells in a batch) (Fig. S2A). The interaction topics represent cell pair interactions across different data sets (Fig. S2B).

##### Topic-specific gene co-expression network

For each interaction topic *t* and LR gene *g*, we calculated a z-score *s*_*tg*_ as 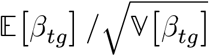, where *β*_*tg*_ is topic-specific gene frequency estimated by the interaction topic model. The highest proportion estimate was used to assign an interaction topic to all cancer cell pairs. A gene co-expression network for each interaction topic was constructed using significantly enriched LR genes (z-score > 4). The edge in the network represents the magnitude of Pearson correlation coefficient between LR gene pairs based on expression data in all the cancer cell pairs assigned to respective interaction topic.

## STATISTICAL ANALYSIS

We performed deep learning using `PyTorch` library (https://pytorch.org/) as specified in the above section. We implemented custom-built visualization scripts in `R` language.

## ADDITIONAL RESOURCES

We shared the full working directory of model estimation and statistical analysis via the public repository.

