## Supplemental for "Single-cell Pairwise Relationships Untangled by Composite Embedding model"

**Figure S1. Sensitivity analysis and model estimation traces of cell topic models**

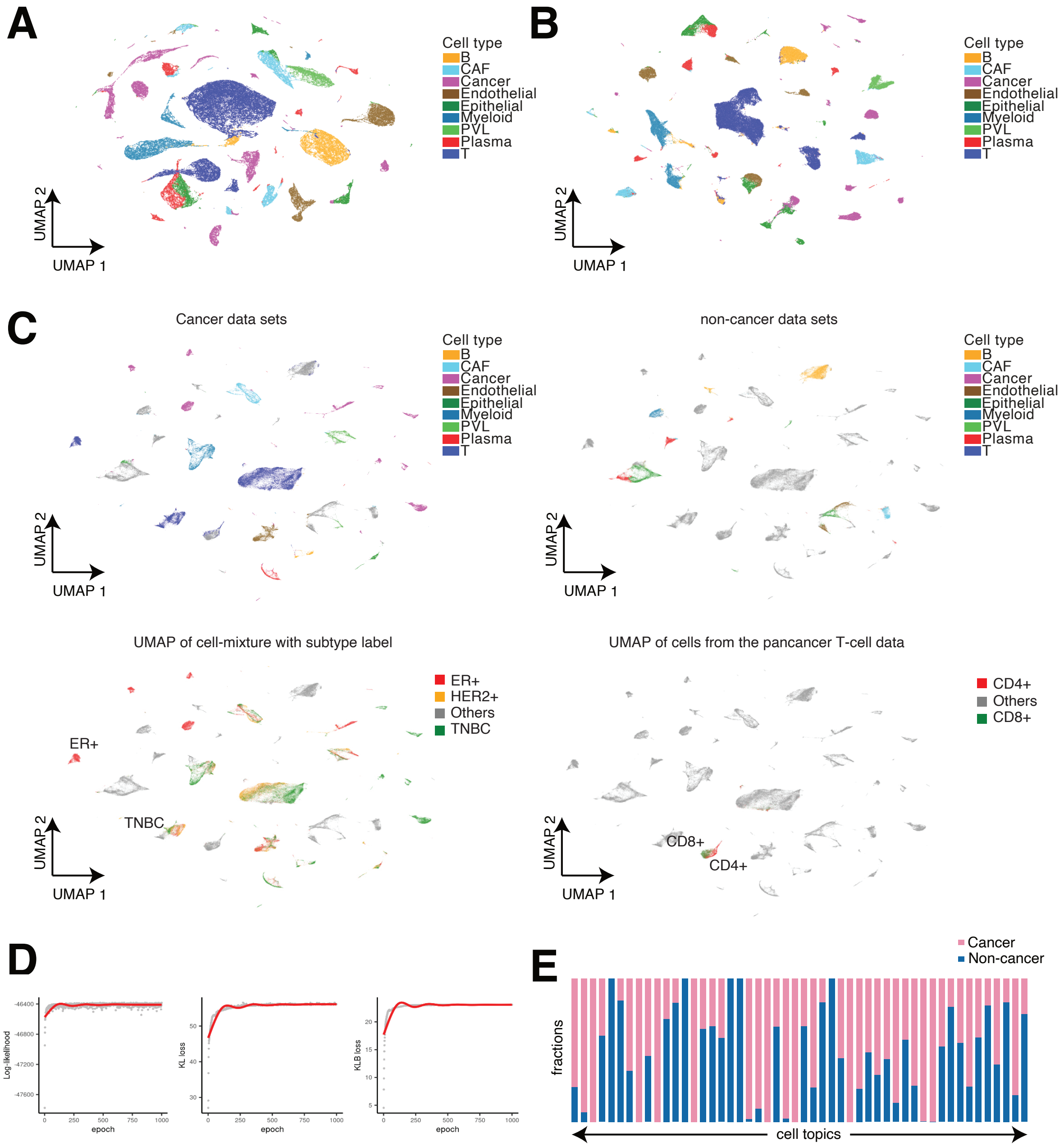


**(A)** UMAP visualization of cell topics (k=10). **(B)** UMAP visualization of cell topics (k=25). (C)UMAP visualization of cell topics (k=50) from different data sources - breast cancer data, normal breast data, immune cells from pan cancer data, and breast cancer subtypes. (D) The model training trace for cell topic model (k=50). (E) The distribution of proportion of cells in cell topics from cancer vs. normal data sources.

### Figure S2. Model estimation traces for Interaction topic model

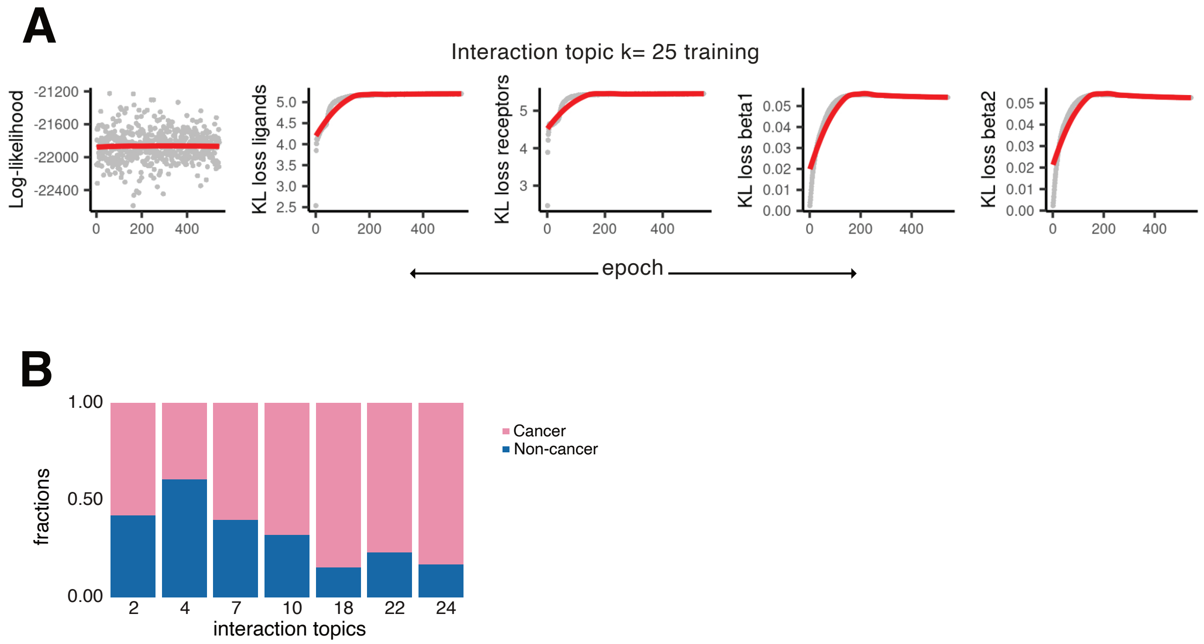


**(A)** Log-likelihood and KL loss trace in training for the interaction topic model.

**(B)** The fraction of cancer/non-cancer cells within the seven interaction topics.

### Figure S3. Overlap with existing interaction databases

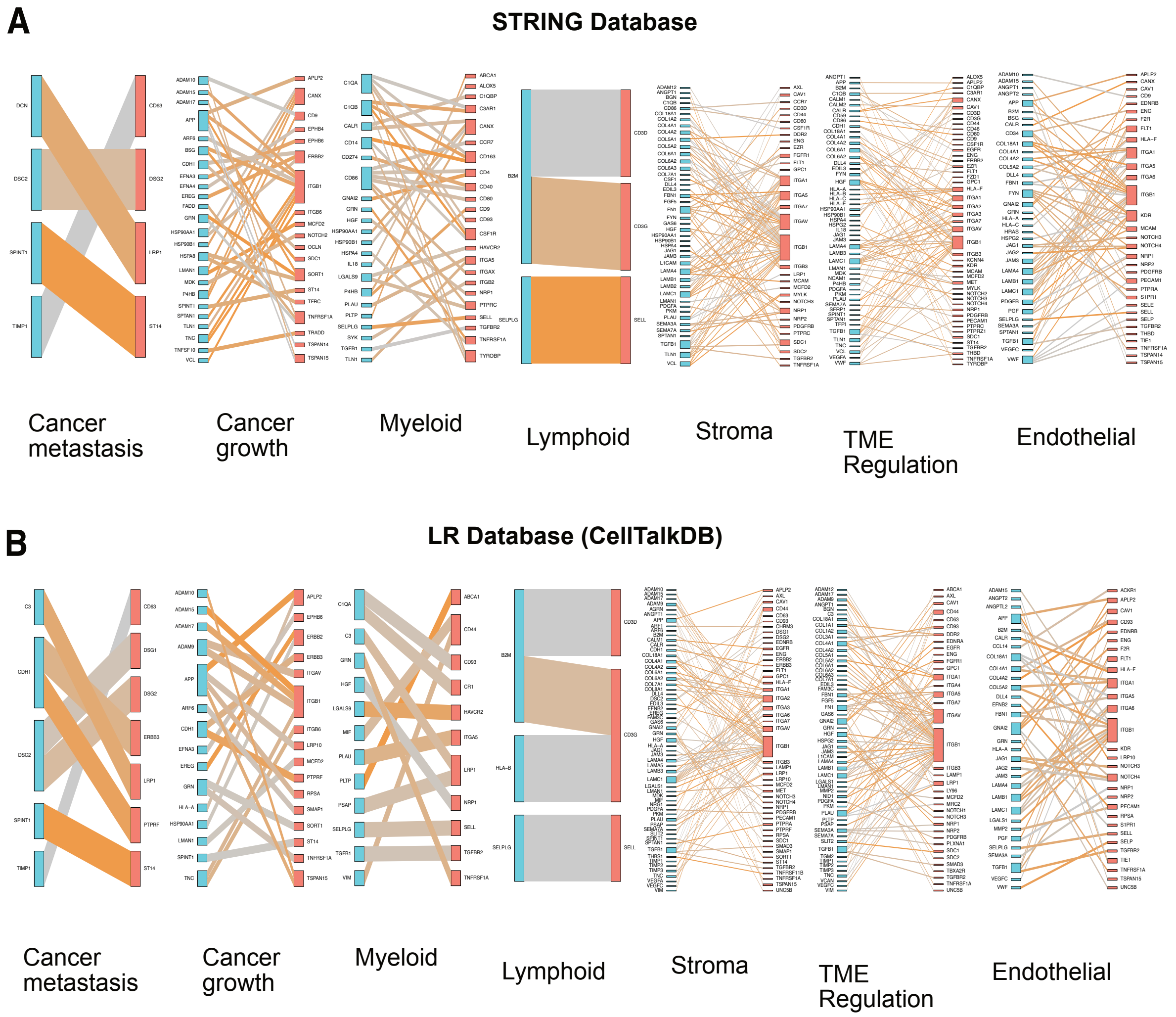


Overlap of gene-gene interactions derived from iteration topic model with other gene networks in **(A)** STRING database (v11.5 with combined score > 0.3) (Szklarczyk et al. 2019) and **(B)** CellTalk database (v20220131) (Shao et al. 2021).

Table 1:

| Topic | Size (#pairs) | Percent | Cell type | Top Ligand Genes | Top Receptor Genes |
|---|---|---|---|---|---|
| 0 | 596,396 | 2.41 | Cancer | GDF5, NTNG1, SPINK1, FGF5, BCAN | EPHA5, GPR182, STAB2, PLXNB3, LRRC4C |
| 1 | 606,439 | 2.45 | Cancer | GDF5, NTNG1, FGF5, BCAN, SPINK1 | EPHA5, GPR182, STAB2, PLXNB3, LRRC4C |
| 2 | 2,134,165 | 8.61 | Endothelial | ANGPT2, BSG, CCL14, CD34, NID2 | APLNR, BOC, CD93, ESAM, SELE |
| 3 | 583,837 | 2.36 | Cancer | GDF5, NTNG1, SPINK1, FGF5, BCAN | EPHA5, GPR182, STAB2, GRIN2B, PLXNB3 |
| 4 | 858,872 | 3.46 | Epithelial | GRP, KISS1, NLGN3, SCGB3A2, TPH1 | IL20RB, KISS1R, KLB, MMP24, SLC17A7 |
| 5 | 613,330 | 2.47 | Cancer | GDF5, NTNG1, SPINK1, FGF5, BCAN | EPHA5, GPR182, STAB2, PLXNB3, GRIN2B |
| 6 | 576,325 | 2.32 | Cancer | GDF5, SPINK1, NTNG1, BCAN, FGF5 | EPHA5, GPR182, STAB2, PLXNB3, GRIN2B |
| 7 | 1,200,434 | 4.84 | Epithelial | DLK2, EDN3, GREM1, NMB, NTF3 | COL17A1, GRPR, IL1RL1, NTRK3, UNC5A |
| 8 | 588,685 | 2.37 | Cancer | GDF5, NTNG1, SPINK1, BCAN, FGF5 | EPHA5, GPR182, STAB2, PLXNB3, GRIN2B |
| 9 | 613,759 | 2.48 | Cancer | GDF5, SPINK1, NTNG1, FGF5, BCAN | EPHA5, GPR182, STAB2, PLXNB3, LRRC4C |
| 10 | 2,265,596 | 9.14 | Cancer | C4B, HSP90AA1, LEFTY1, SLIT1, UCN | ADORA2A, ERBB2, FZD10, PLA2R1, TSPAN1 |
| 11 | 622,494 | 2.51 | Cancer | GDF5, NTNG1, SPINK1, BCAN, FGF5 | EPHA5, GPR182, STAB2, PLXNB3, LRRC4C |
| 12 | 626,524 | 2.53 | Cancer | GDF5, NTNG1, SPINK1, BCAN, FGF5 | EPHA5, GPR182, STAB2, PLXNB3, LRRC4C |
| 13 | 629,455 | 2.54 | Cancer | GDF5, NTNG1, SPINK1, BCAN, FGF5 | EPHA5, GPR182, STAB2, PLXNB3, LRRC4C |
| 14 | 630,346 | 2.54 | Cancer | GDF5, NTNG1, SPINK1, BCAN, FGF5 | EPHA5, GPR182, STAB2, LRRC4C, PLXNB3 |
| 15 | 607,261 | 2.45 | Cancer | GDF5, NTNG1, SPINK1, BCAN, FGF5 | EPHA5, GPR182, STAB2, PLXNB3, GRIN2B |
| 16 | 633,812 | 2.56 | Cancer | GDF5, NTNG1, SPINK1, FGF5, BCAN | EPHA5, GPR182, STAB2, PLXNB3, LRRC4C |
| 17 | 590,853 | 2.38 | Cancer | GDF5, NTNG1, BCAN, SPINK1, FGF5 | EPHA5, GPR182, STAB2, PLXNB3, LRRC4C |
| 18 | 2,269,902 | 9.16 | CAF | COL1A1, COL1A2, COL3A1, MMP13, NID2 | EPHA5, GPR1, ITGA11, KCND2, SCARA5 |
| 19 | 641,247 | 2.59 | Cancer | GDF5, NTNG1, SPINK1, FGF5, BCAN | EPHA5, GPR182, STAB2, PLXNB3, SLC4A11 |
| 20 | 616,550 | 2.49 | Cancer | GDF5, NTNG1, SPINK1, BCAN, FGF5 | EPHA5, GPR182, STAB2, PLXNB3, LRRC4C |
| 21 | 579,652 | 2.34 | Cancer | GDF5, SPINK1, NTNG1, BCAN, FGF5 | EPHA5, GPR182, STAB2, PLXNB3, GRIN2B |
| 22 | 3,044,872 | 12.28 | T | CLEC2B, CLEC2D, HLA-E, MICB, ULBP2 | CD3D, KIR3DL1, KLRC1, KLRC2, KLRD1 |
| 23 | 579,346 | 2.34 | Cancer | GDF5, NTNG1, SPINK1, BCAN, FGF5 | EPHA5, GPR182, STAB2, PLXNB3, LRRC4C |
| 24 | 2,080,015 | 8.39 | Myeloid | APOC2, C1QA, CCL13, CCL18, CCL23 | CD68, CR1, KCNJ10, LILRB2, TREM2 |

### Figure S4. Annotations for 25 Interaction Topics
